# Colonies of *Acropora formosa* with greater survival potential show conservative calcification rates

**DOI:** 10.1101/2020.09.28.315788

**Authors:** Vanessa Clark, Matheus A. Mellow-Athayde, Sophie Dove

## Abstract

Coral reefs are facing increasingly devasting impacts from ocean warming and acidification due to anthropogenic climate change. In addition to reducing greenhouse gas emissions, potential solutions have focused either on reducing light stress during heating, or the potential for identifying or engineering “super corals”. These studies, however, have tended to focus primarily on the bleaching response of corals, and assume that corals that bleach earlier in a thermal event are more likely to die. Here, we explore how survival, potential bleaching, and coral skeletal growth (as branch extension and densification) varies for conspecifics collected from distinctive reef zones at Heron Island on the Southern Great Barrier Reef. A series of reciprocal transplantation experiments were undertaken using the dominant reef building coral (*Acropora formosa*) between the highly variable ‘reef flat’ and the less variable ‘reef slope’ environments. Coral colonies originating from the reef flat had higher rates of survival and thicker tissues but reduced rates of calcification than conspecifics originating from the reef slope. The energetics of both populations however benefited from greater light intensity offered in the shallows. Reef flat origin corals moved to the lower light intensity of reef slope reduced protein density and calcification rates. For *A. formosa*, genetic difference, or long-term entrainment to a highly variable environment, appeared to promote coral survival at the expense of calcification. The response divorces coral resilience from carbonate coral reef resilience, a response that was further exacerbated by reductions in irradiance. As we begin to discuss interventions necessitated by the CO_2_ that has already been released to the atmosphere, we need to prioritise our focus on the properties that maintain valuable carbonate ecosystems. Rapid and dense calcification by corals such as branching *Acropora* is essential to the ability of carbonate coral reefs to rebound following disturbances events, but may be the first property that is sacrificed to enable coral genet survival under stress.

## Introduction

Primary (skeletal expansion) and secondary (skeletal densification) calcification by reef-building corals is critical to reef construction and other services provided by coral reef ecosystems (Spurgeon 1992). Environmental impacts on rates of net coral calcification often differentiate ecosystems that are labelled carbonate coral reef ecosystems and those that are labelled coral reef communities (Kleypas et al. 1999a). The ephemeral nature of reef-building by Scleractinian corals has, however, enabled corals to survive mass extinction events (Stanley 2006). Calcification is energy expensive (Cohen and Holcomb 2009), and in some environments, it may be necessary to redirect energy investment away from calcification in order to facilitate coral survival (Leuzinger et al. 2012). Here, we evaluate drivers such as putative epigenetic responses and exposure to reduced light levels over summer and their influence on primary and secondary calcification for the branching coral, *Acropora formosa* (Dana, 1846). *Acropora formosa* is present in diverse zones of a coral reef where it contributes to the 3-dimensional structure and carbonate accumulation within coral reef ecosystems (Veron 1993).

Underwater marine heat waves are increasing globally and are impacting multiple ecosystems including the Great Barrier Reef (Hoegh-Guldberg 1999; Hughes et al. 2018a; Smale et al. 2019). Initially, coral bleaching observed over a large area drew our attention to the fact that ocean warming was negatively affecting coral hosts and their endosymbiotic dinoflagellates (Hoegh Guldberg and Salvat 1995). More frequent and intense thermal heatwaves are now associated with mass coral mortality, where corals die irrespective of their apparent bleaching susceptibilities (Hughes et al. 2018b; Dove et al. 2020). Past emission of CO_2_ from fossil fuel burning, and the lag between emission and ocean impacts, mean that future reefs will almost certainly be exposed to thermal events that will occur with greater frequency and greater intensity than those we have witness to date (Hoegh-Guldberg et al. 2019). The adoption of CO_2_ mitigation pathways such as that agreed in Paris at the United Nations Climate Change Conference in 2015 will significantly reduce ocean acidification and hence impacts on coral reef ecosystems (Magnan et al. 2016). Eventuating conditions are nonetheless likely to have impacts on coral communities (Albright et al. 2016). Impacts that may affect the rate of net calcification and frame work-building on a coral reef, thereby threatening key services provided by these ecosystems (Hoegh-Guldberg et al. 2007).

The degree to which these ecosystems are threatened under reduced CO_2_ emission scenarios such as RCP2.6 (IPCC 2014) is nonetheless debated with some arguing that epigenetics can accelerate coral adaptation either naturally (Palumbi et al. 2014) or via assisted evolution (Van Oppen et al. 2015; Putnam et al. 2016). In such studies, bleaching susceptibility is often viewed as interchangeable with mortality susceptibility (van Oppen et al. 2011; Puill-Stephan et al. 2012; Levin et al. 2017; Cleves et al. 2018), and the costs (in terms of skeletogenesis) associated with “adaptation” are rarely considered (Hoegh-Guldberg et al. 2011; Pandolfi et al. 2011). Other research efforts have focused on potential solutions associated with reducing light stress to inhibit thermal bleaching (Jones et al. 2016). Such potential solutions assume corals and their endosymbionts do not acclimate sufficiently rapidly to reductions in light to obviate the benefits of reducing light on symbiont excitation pressure at PSII reaction centres (Iglesias-Prieto and Trench 1997), with excessive excitation driving the formation of reactive oxygen species that can trigger a bleaching response in corals (Smith et al. 2005; Dove et al. 2006). At the same time, they assume that the reduction in light and hence reduced photosynthesis from periods of shading is not important to host coral energetics (Fisher et al. 2019). In the present study, we examined the potential for either of these solutions to translate into increased survival potential and optimal framework building in response to summer conditions (2017-2018). We also argue that, if they are not beneficial under current conditions, then they are unlikely to be beneficial under future climate conditions.

The study was done at two different reef locations, (a) the shallow reef flat (1-3 m) and (b) the deeper reef slope (7-8 m) of Heron Island, on the Great Barrier Reef. Colonies from the reef flat sites experienced a large diel range and variability in sea water chemistry (e.g. pCO_2_), temperature and light intensity, which is associated with the large tidal range (3m) and the ponding of reefs at low tide (McGowan et al. 2010; Santos et al. 2011; Kline et al. 2012, 2015). Periods of excessively high (in summer) or low (in winter) temperatures associated with ponding can inhibit the efficiency of photosynthesis (Coles and Jokiel 1977) given Heron Island’s high latitude location. Over such periods, corals growing at Heron Island potentially limited in their heterotrophic uptake capacity may be forced to consume previously deposited biomass to facilitate genet survival (Anthony et al. 2009; Grottoli and Rodrigues 2011). Survival in such environments may therefore favour organisms with larger biomass per unit surface area (Szmant and Gassman 1990; Grottoli et al. 2006), a property that in turn may be facilitated by reductions in coral expansion and/or in rates of secondary calcification (Anthony et al. 2002; Tanaka et al. 2007).

By contrast, on the reef slope, temperature and pCO_2_ conditions are relatively stable due its proximity and exchange with external ocean waters (Albright et al. 2015). Light, however, attenuates significantly with depth and can constrain phototrophy. On the other hand, the increased exposure to high energy of waves tends to limit the potential for net colony expansion, especially for corals that deposit less dense skeletons over the long term (Allemand et al. 2011). The combination of fragmentation, driven by high wave energy and rapid rates of skeletogenesis can spread the risk of genotype mortality across a larger area (Highsmith 1982). Therefore, it may be hypothesized that, in general, fast growing morphotypes are more frequently associated with high energy locations; whilst fleshier morphotypes, that tend to invest more in tissue than skeletal expansion, are associated with more marginal, low energy habitats (Hoogenboom et al. 2008).

Epigenetic memory potentially offers corals and other organisms a mechanism for responding to climate change at rates that are greater than one might expect based on their average generation times (Van Oppen et al. 2015; Putnam et al. 2016 vs Hoegh-Guldberg et al. 2002). Epigenetic memory is a mechanism through which the transcriptional responses of parent cells to environmental stimuli are maintained as the favoured set of transcriptional responses in descendant cells irrespective of environmental stimuli (Klosin et al. 2017). In this sense, epigenetic exposure to highly variable light and/or temperature environments, can lead to transcriptional behaviours in parent cells that are retained in descendant cells. Descendant cells are therefore less likely to be responsive to their immediate environment, but rather retain the properties and behaviours that enabled parent cells to survive and divide in their relatively hostile historic environment. The long-term prior experience of a coral genet that settled and grew to adulthood in a highly variable environment, does, in at least some cases, appear to provide an epigenetic entrainment that increases the survival potential of corals experimentally exposed to regionally anomalous increases in temperature (Oliver and Palumbi 2011; Grottoli et al. 2017; Thomas et al. 2018). Some such corals have been referred to as “super corals” (Grottoli et al. 2017; Darling and Côté 2018). This term has been used in reference to naturally occurring robust species of corals, conspecifics that have settled and developed into adult colonies in hotter and/or more variable environments (Oliver and Palumbi 2011; Grottoli et al. 2014, 2017), but also to describe the artificial manipulation of corals through the formation of chimeras (van Oppen et al. 2011; Puill-Stephan et al. 2012), editing of larval genes (Cleves et al. 2018), or the insertion of artificially developed symbionts (Levin et al. 2017). “Super corals” are first and foremost described as having a higher thermal tolerance in the face of bleaching, requiring a more extreme temperature to induce the loss of symbionts (Hoogenboom et al. 2008; Grottoli et al. 2017). The cost of this bleaching resilience, however, in terms of both aspects of skeletogenesis has not been explored. Here, the link between bleaching sensitivity and greater host survival is often assumed, along with the assumption that corals conspecifics that do not bleach are healthier than those that do bleach (Berkelmans and Van Oppen 2006; Howells et al. 2013; Lirman et al. 2014).

Implicit to coral health is the assumption that their endosymbiotic dinoflagellates are strict photo-autotrophic obligate mutualists, which are incapable of translocating organic carbon from host to symbiont (Rowan and Knowlton 1995; Rowan et al. 1997; Mieog et al. 2009). Relatively recent advances in the literature, however, indicate that this is incorrect as all chloroplast-containing dinoflagellates, inclusive of Symbiodiniaceae, are facultative heterotrophs (Steen 1987; Mulholland et al. 2018). At least some are capable of phagotrophy, but all are capable of osmotrophy (Flynn et al. 2013). And hence, all, under the appropriate changes to the environment, can become parasitic on their coral hosts. Several studies clearly demonstrate that non-bleached corals can be even less healthy than their unhealthy bleached counterparts. Non-bleached conspecifics can show higher susceptibility to polyp mortality when phototrophic capacity is eliminated (Steen 1986, 1987). Similarly, bleached hosts that house thermally tolerant symbionts can show reduced extension rates, lower lipid densities, and decreased reproductive output both during and after the advent of a thermal event (Jones and Berkelmans 2010, 2011). The visual nature of mass coral bleaching has led us to overly focus on the symbiont response, as opposed to realising that the host appears to be energetically deprived when exposed to thermal stress often irrespective of whether symbiont population densities are maintained. Recent observations of prolonged thermal events that lead to mass coral mortality irrespective of the documented bleaching susceptibility of the taxa involved (Hughes et al. 2018b; Dove et al. 2020) would also seem to argue for a new functional definition for a “super coral”. One that focusses directly on survival and growth of corals as opposed to their brown colour and bleaching sensitivity.

We ran reciprocal field experiments on *Acropora formosa* growing in two distinct environments; on the highly variable shallow reef flat and in a less variable, but slightly deeper outer reef slope of Heron Island, on the southern Great Barrier Reef, Australia. We aimed to test an alternative hypothesis that corals survive highly variable environments because they engage in conservative extension rates that enable them to invest their energy into tissue maintenance and production. We further aimed to explore whether there was any influence of prior environmental history (i.e. potentially epigenetic memory) that might constrain the responses of colonies of *A. formosa*. Specifically, we sought to: (i) determine whether higher exposure to variability in prior life history increases coral survival irrespective of the characteristic of the new environment; (ii) Whether properties that potentially increase survival are traded for reduced rates of calcification; (iii) whether light reductions associated with alteration in depth and reef location, led to a rapid acclimation response or not and whether a potential decrease in energy acquisition occurs as corals face changed environmental conditions.

## Materials and Methods

### Study Site

Heron Reef (23°27’S, 151°55’E) is a lagoonal platform reef which is part of the Capricorn Bunker group located on the southern end of the Great Barrier Reef, Australia. Past sea surface temperatures (SST) for the sea surrounding Heron Reef had a maximum monthly mean (MMM) of 27.3°C (between 1985 and 2001; Weeks et al. 2008). This is also the MMM used to determine hotspots and degree heating weeks (DHW), and thereby exposure to thermal stress (Liu et al. 2014). Heron Reef has a semi-diurnal tide cycle, which drives a large scale variation in temperature, oxygen, and carbon dioxide (Kline et al. 2012). For this study, sites were either located in reef flat or outer reef slope on the southern side of Heron Island tides. Conditions on the reef flat (1-3m, Gourlay and Jell 1992) are highly variable with regular periods of ponding with minimal or no inward or outward exchange of water (Kinsey 1967; Jell and Flood 1978; Potts and Swart 1984; Jell and Webb 2012; Georgiou et al. 2015). The reef slope site (depth 7-8m at mid-tide, Harry’s Bommie) was on the outer margin of the reef nearby Wistari channel and hence exposed to well mixed ocean waters along with significant wind and wave activity (Bradbury 1981; Connell et al. 1997). Brown et al. (2018) conducted studies at similar sites on Heron Island finding that mean photosynthetically active radiation (PAR) was two to three times higher on the reef flat than the reef slope throughout the year, while mean temperature remained relatively similar between sites seasonally. Heron Reef is dominated by *Acropora spp*. (Santos et al. 2011; Brown et al. 2018) which are typically fast growing under normal conditions, and as a result can rapidly recolonise after events that have led to reef degradation (Highsmith 1982), especially via fragmentation. Making them fundamental in maintaining positive carbonate balances within coral reef communities (Brown et al. 2020).

### Reciprocal Transplantation

Fragments (7-10cm) of *A. formosa* were collected from different colonies at each location (‘Flat’ n=125; ‘Slope’ n=125) in late September 2017. Fragments were randomly collected from the growing tips of colonies, fragments with axial branches or damaged apical corallites were excluded. While these were selected from different colonies to avoid pseudo-replication it was not tracked. Corals were kept in flow through aquaria under a shade cloth during initial measurements and preparation for the different experimental treatments. All fragments were trimmed to 7cm using bone cutter scissors. A total of 25 fragments from each location (Flat, Slope) were selected to represent initial measurements, theses were stored at - 20°C until being analysed either at Heron Island Research Station and/or the University of Queensland. The remaining fragments had a hole drilled ∼1cm from the base of the nubbin for ease of attachment to live rock. Ties through the hole were used to stabilise coral nubbins and reduce algal growth (see fig 2). Nubbins were randomly assigned to treatments in a fully crossed design (Fig 1), corals originating from the reef flat were either returned to the reef flat (“Origin[Flat]/ Transplant[Shallow]”, n=50), or transplanted to the reef slope (“Origin[Flat/ Transplant[Deep]”, n=50), and corals originating from the reef slope were either returned to the reef slope (“Origin[Slope]/ Transplant[Deep]”, n=50), or transplanted to the reef flat (“Origin[Slope]/ Transplant[Shallow]”, n=50). Once attached to live rock corals were deployed on metal frames at each location in late September 2017. The live rocks with nubbins attached were collected in late November 2017 for observation, all dead corals were removed (n=24) prior to redeployment. Final collection of the fragments was conducted in mid-February 2018, each nubbin had final buoyant weight measurements taken before being stored at −20°C until further analysis could be undertaken. Specific methodologies for individual measurements are described below. All physiological measurements, apart from buoyant weight, were reported as total change over the experimental period based on the average value calculated from the 25 fragments collected and processed at the beginning of the experiment.

**Fig 1.**
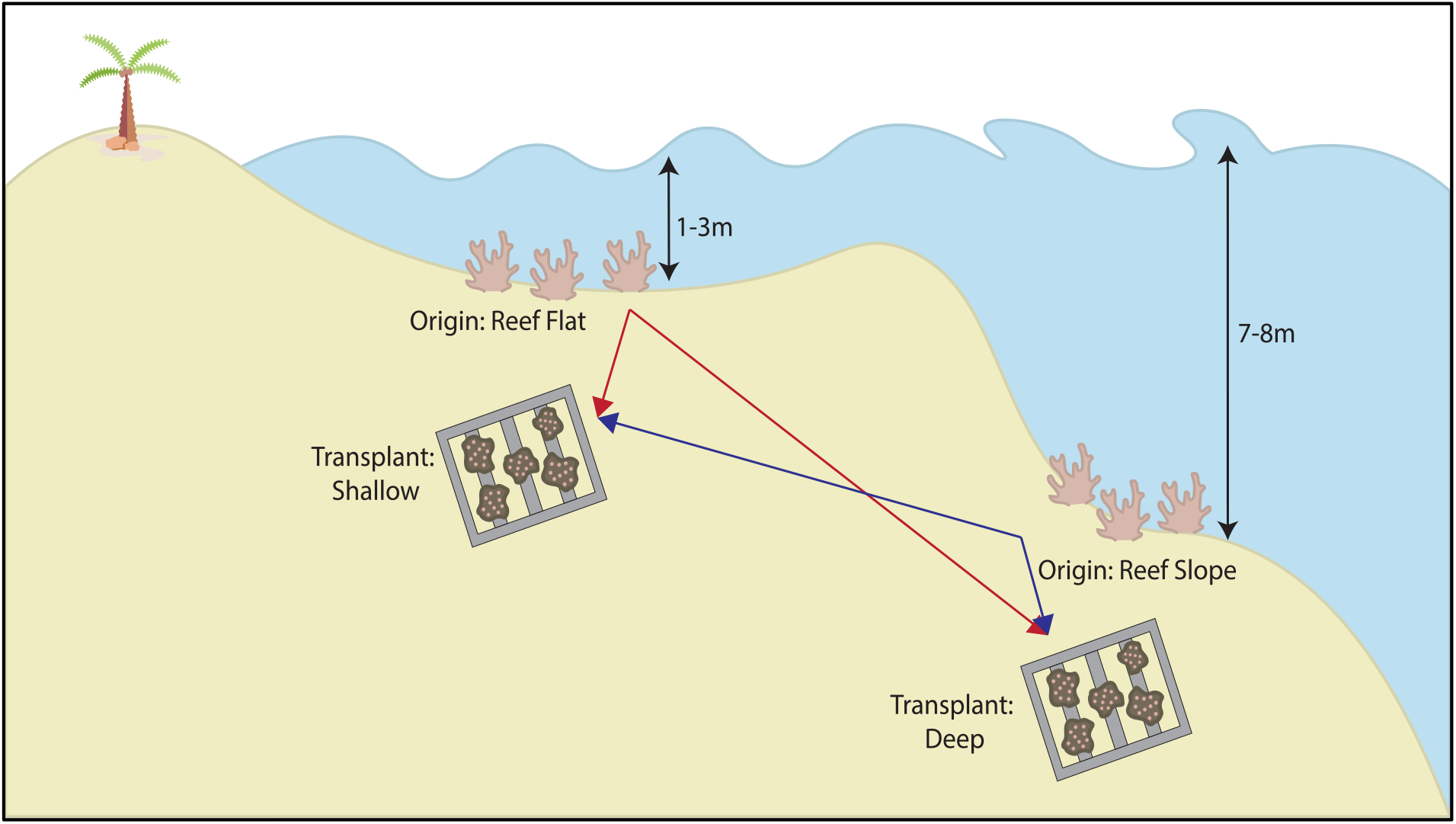
Coral fragments were transplanted between locations on the reef flat and reef slope of Heron Island in the Southern Great Barrier Reef of Australia. For clarity, sites of colony Origin are referred to as “reef slope” and “reef flat”, and locations of experimental positioning (transplantation) are referred to as “deep” and “shallow”. Created in Adobe Illustrator.

**Fig 2.**
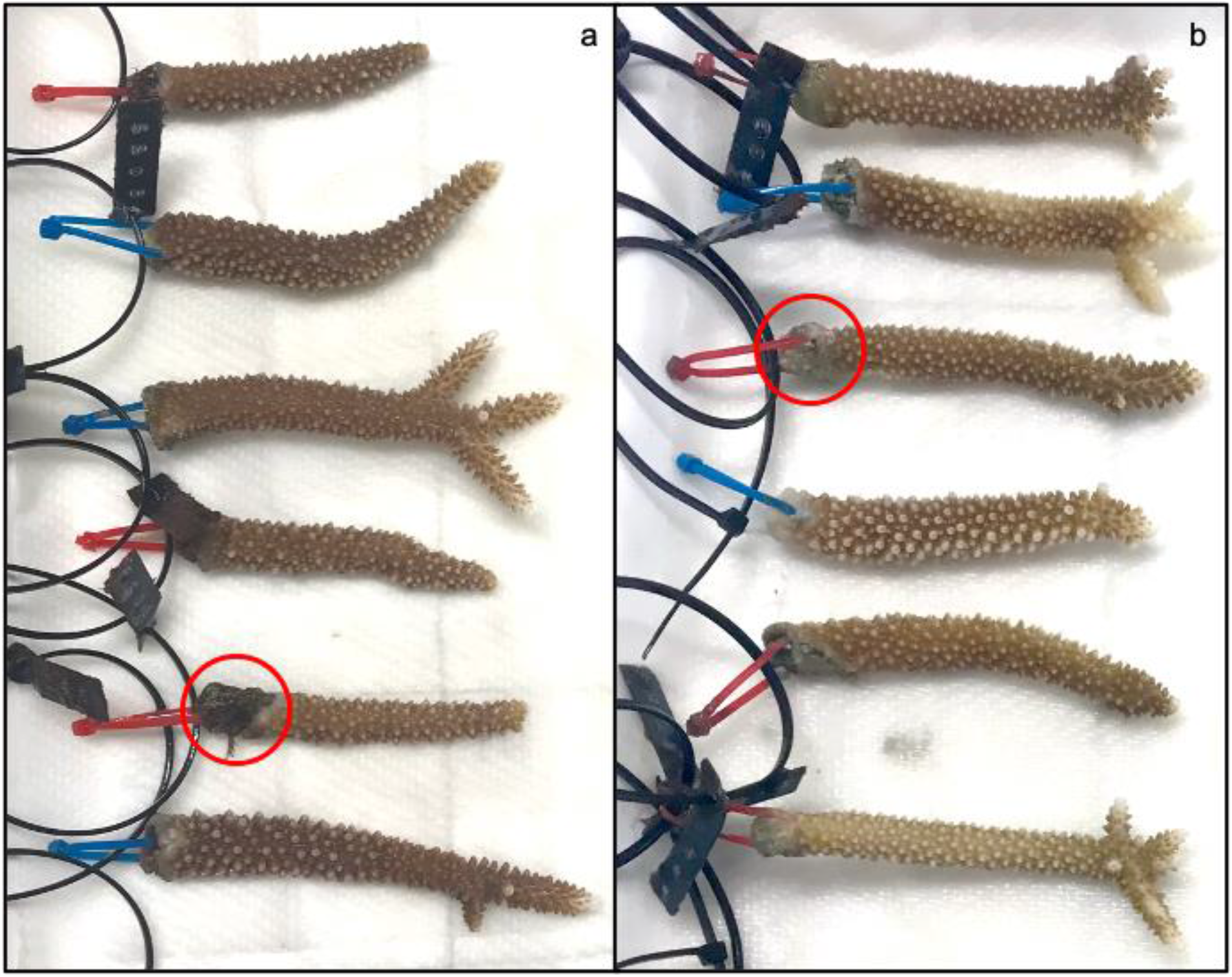
End-point examples of *Acropora formosa* nubbins following a reciprocal transplantation experiment conducted at Heron Island (Southern GBR). (a) Nubbins experimentally located as transplants to the “deep” reef slope, and (b) Nubbins experimentally located as transplants to the “shallow” reef flat. Zip-ties represent location of colony Origin (slope = blue, flat = red). Tissue dieback can be observed at the base of nubbins where zip-ties have been attached, differential levels of dieback are due to a lack of uniformity in the live rock corals were attached to. The red circle in (a) highlights dieback with algal growth, and in (b) highlights dieback with the absence of coralites and tissue.

### Physiological analyses

Calcification rates were calculated from initial and final buoyant weight measurements of each individual fragment. Buoyant weight was measured using methods described by Spencer Davies (1989) and Jokiel et al. (2008). Coral tissue was removed from the skeleton by airbrush using 0.06 M phosphate buffer solution (PBS). Skeletons were then cleaned using a 10% bleach solution to remove any residual tissue. If corals had any tissue die-back this was marked on skeletons in preparation for living surface area analysis. Fragments were rinsed and dried at 70°C for two hours. Methods described in Bucher et al. (1998) were used for determining bulk volume.

The modified double wax dipping method described by Holmes (2008) was used to measure the surface area of living tissue. Paraffin wax was heated to 65°C, and coral fragments were maintained at room temperature throughout. This method has been shown to give the most accurate surface area estimation compared to X-ray CT scanners for Acroporid corals (Naumann et al. 2009; Veal et al. 2010).

The coral tissue in PBS was centrifuged at 4500rpm for 5 minutes (3K15 Sigma laborzentrifugen GmbH, Osterode, Germany), to separate the symbiont pellet. The resulting supernatant was stored in two parts to allow for protein and lipid analyses. For protein analyses the supernatant was analysed with a spectrophotometer (Spectramax M2 Molecular Devices, California, USA), using absorbance values at 235nm and 280nm. Protein concentrations were then calculated using equations in Whitaker and Granum (1980). For lipid analyses, the supernatant was stored at −80°C overnight. Frozen samples were placed in a freeze dryer (Coolsafe 9l Freeze Dryer Labogene, Allerød, Denmark) at −110°C for 24 to 48 hours until all moisture was removed. Total lipid extraction from each sample was conducted using methods described in Dunn et al. (2012).

### Mortality

Fragments in the reciprocal transplant could only be identified as dead in collection periods, at which point they were removed from the live rock. At the midway point of the study, 24 corals were classified as dead and removed, with one additional coral removed due to mortality at the end point of the study. A coral fragment was classified as dead if it lacked coral tissue and had been colonised by algae.

Tissue dieback on all live fragments was also assessed at the end of the experiment. Approximately 80% of shallow transplants, and 77% of deep transplant experienced a small amount of dieback near the point of attachment to the live rock. Algal growth was also apparent in some cases (fig 2a), and the absence of coralites and tissue in others (fig 2b). This was marked and accounted for when conducting live surface area measurements. Additionally, 5% of deep transplants exhibited tissue dieback beyond the attachment region but not total mortality, these were excluded from the analysis of physiological properties.

### Environmental Variables

Field environmental variables were measured using temperature (Hobo pendent logger Onset, MA, USA), and photosynthetically active radiation (PAR; Odyssey logger Dataflow systems LTD, Christchurch, NZ) loggers. Three PAR and three temperature loggers were placed at each location (“deep” reef slope, and “Shallow” reef flat) each within 5 metres of the metal frames on which the corals were attached. On the reef slope, two of the three PAR loggers failed and as a result only one set of data was able to be used for analyses. Temperature loggers collected data as hourly means; PAR loggers collected data as integrated values every 2 hours. For analyses, 24h daily means, 25^th^ percentile (Q1) and 75^th^ percentile (Q3) were determined for temperature, but 12h (6h – 18h) daytime mean, Q1 and Q3 were determined for PAR.

### Statistical Analyses

All statistical analyses were conducted with R version 3.6.2 software (2018), figures were created using the ggplot2 package (Wickham 2016) and sjPlot package (Lüdecke 2014). Data was assessed for normality and homogeneity of variance through visual inspection of Q-Q plots and residual vs fitted values prior to analysis. All physiological data (bulk volume (BV), living surface area (LSA), buoyant weight (BW), total protein, and total lipids) was initially calculated as total change over time, however after comparison of initial bulk volume between locations of Origin, this was converted to percentage change. This meant that final reported values accounted for the basal variability in bulk volume between fragment dependent on their location of Origin. Linear mixed effects (LME) models were then developed using a stepwise procedure, Akaike information criterion was applied in each case to select the model with the best fit. Models with different response variables were built up in order of increasing complexity in order to explore trades between variables (Table 1). The function Anova (John et al. 2020) was applied to models, with the type 2 or 3 selection of treatment of sum of squares based on whether optimal models suggested significant interactions amongst variables.

**Table 1.**
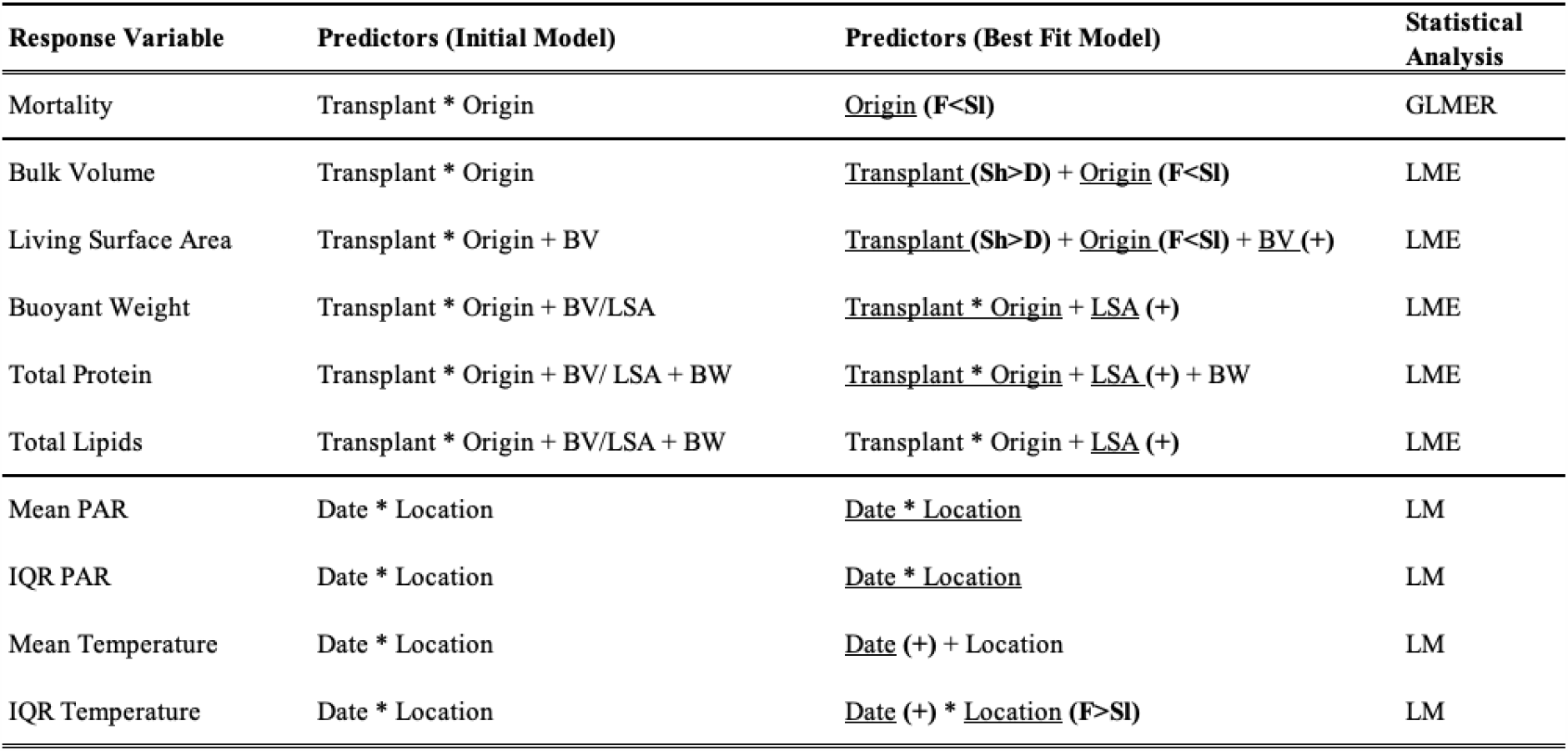
Response and predictor variables for the initial and best fit models applied to the analysis of samples from the reciprocal transplantation experiment conducted on Heron Island (Southern GBR). Underlined predictor in best fit model indicate significant effect (P<0.05). The direction of these significant effects is indicated in brackets, for continuous predictors (+) defines a positive correlation; for categorical predictors, the relationship between levels of the factor are provided. Transplant, Sh = Shallows, D = Deep; Origin, F = Flat, and Sl = Slope. The direction of interaction effects is not described in this table. Variables are abbreviated in model outputs; bulk volume = BV, living surface area = LSA, buoyant weight = BW.

| Response Variable | Predictors (Initial Model) | Predictors (Best Fit Model) | Statistical Analysis |
| --- | --- | --- | --- |
| Mortality | Transplant * Origin | <u>Origin</u> (F<SI) | GLMER |
| Bulk Volume | Transplant * Origin | <u>Transplant</u> (Sh>D) + <u>Origin</u> (F<SI) | LME |
| Living Surface Area | Transplant * Origin + BV | <u>Transplant</u> (Sh>D) + <u>Origin</u> (F<SI) + <u>BV</u> (+) | LME |
| Buoyant Weight | Transplant * Origin + BV/LSA | <u>Transplant</u> * <u>Origin</u> + <u>LSA</u> (+) | LME |
| Total Protein | Transplant * Origin + BV/LSA + BW | <u>Transplant</u> * <u>Origin</u> + <u>LSA</u> (+) + BW | LME |
| Total Lipids | Transplant * Origin + BV/LSA + BW | Transplant * Origin + <u>LSA</u> (+) | LME |
| Mean PAR | Date * Location | <u>Date</u> * <u>Location</u> | LM |
| IQR PAR | Date * Location | <u>Date</u> * <u>Location</u> | LM |
| Mean Temperature | Date * Location | <u>Date</u> (+) + <u>Location</u> | LM |
| IQR Temperature | Date * Location | <u>Date</u> (+) * <u>Location</u> (F>SI) | LM |

For physiological analysis, model response variables were; BV, LSA, BW, total protein, and total lipids all of which were continuous. Of the fixed predictors, Origin and Transplant were categorical, while the remaining (BV, LSA, and BW) were continuous. Due to collinearity as measured through variance inflation factor (VIF), covariates BV and LSA were identified as interchangeable in subsequent models, the model with the best fit was chosen as previously described. All physiological analysis and mortality models included live rock as a random effect, which refers to the live rock on which the coral was placed, this remained constant throughout the experimental period.

Mortality was analysed as a binomial response variable using a mixed effect logistic regression (GLMER, lme4, Bates et al. 2015) in order to take the random effect of live rock into consideration. Laplacian approximation was used to determine the best fit model. The full and best fit models are shown in Table 1.

A linear model was used to compare environmental variables (Temperature and PAR) between each site (Shallow and Deep) over the four-month experimental period. The daily mean, and interquartile range (IQR, between Q1 and Q3 as previously described), as an indication of variability, were both analysed for temperature and PAR, full and best fit models are shown in Table 1, with date and location as categorical predictors. Degree heating weeks (DHW) were calculated using Q3 to measure heat accumulation over the experimental period, calculations were based on theory from NOAA (Liu et al. 2014) where heating weeks accumulate when temperatures rises above MMM +1.

## Results

### Initial measurements, environmental parameters, and mortality potential

Over the length of the experimental period, corals that originated from the reef slope were 20% less likely to survive than reef flat origin corals (Anova, *χ*^2^ = 4.604, d.f. = 1, *P* = 0.0319, Fig 3a). This occurred over and above the random effect of live rock on mortality (fixed/random effects = 0.04/0.59). *Acropora formosa* fragments from the reef slope had significantly greater initial bulk volume than reef flat origin fragments (Anova, F_(1,46)_=15.42, P=<0.001, Fig 3b).

**Fig 3.**
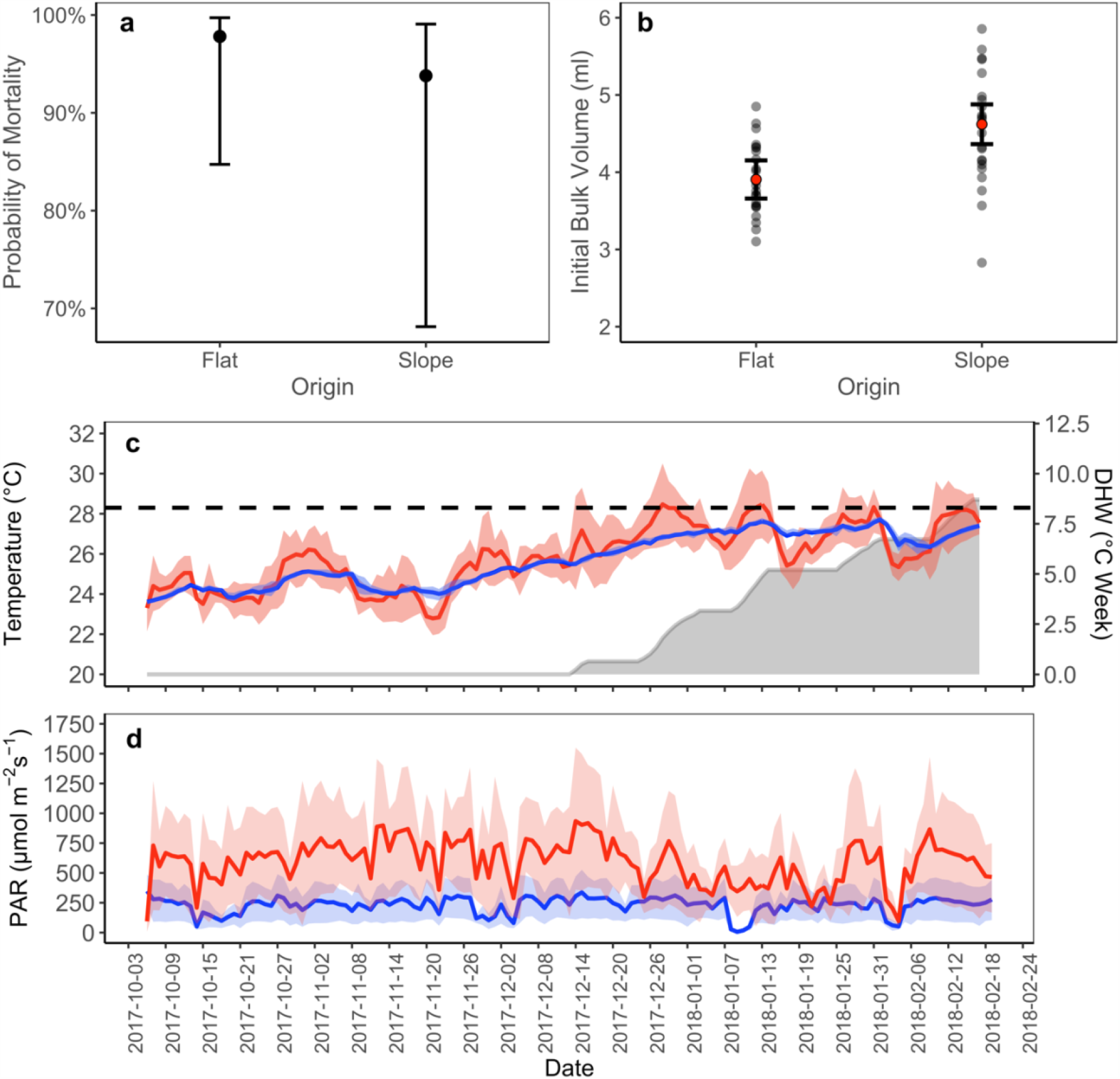
Nubbin mortality potential, initial volume and abiotic conditions associated with a reciprocal transplantation exercise conducted between the reef flat and reef slope of Heron Island (Southern GBR). (a) Probability of fragment mortality by location of Origin. (b) Initial differences in bulk volume (ml) between *A. formosa* fragments from each location of Origin (p<0.05), error bars represent a 95% confidence interval, grey dots show data points, and red dots indicate mean value in data set. (c) *In situ* 24hr mean temperature (°C) and (d) *In situ* daytime mean PAR (µmol m^−2^ s^−1^) between 6:00 h and 18:00 h. For the deep site (depth = 7-8m, blue line), and the shallow site (depth = 1-3m, red line) between the 6/10/2017 and the 17/02/2018. Ribbons on both figures represent 25^th^ percentile (Q1) on the bottom, and 75^th^ percentile (Q3) on the top, of daily data recorded at each location. The dashed line in figure (c) represents the local MMM + 1 (28.3°C) which is associated with bleaching threshold. Grey area under curve indicates Q3 degree heating week (DHW) accumulation on the reef flat.

A significant interactive effect was observed between date and location for mean PAR (Anova, F_(3,270)_= 20.867, P= <0.001, Fig 3d). Mean PAR over the experimental period was always higher in the shallows than the deep. An increase over the experimental period was observed at both locations but was steeper in the shallows. Variability in PAR, measured as the interquartile range, at both locations also differed as an interactive effect of date and location (Anova, F_(3,270)_= 8.852, P= 0.003, Fig 3d). As with mean PAR an increase over the experimental period was observed at both locations but was much greater in the shallows throughout the experimental period.

Mean temperature increased over time at both sites (Anova, F_(3,290)_= 830.855, P=<0.001, Fig 3d) although mean temperature did not differ between the shallow site and deep site. Variability, however, was significantly higher in the shallow than in the deep (Anova, F_(3,290)_= 125.179, P=<0.001, Fig 3d). Temperature variability significantly increased at both locations over the experimental period (Anova, F_(3,290)_=9.205, P=0.003, Fig 3c). The Q3, and mean to a lesser degree, periodically breached the MMM+1 (28.3°C) threshold established for the greater region, in the shallow but not in the deep. Q3 of the shallow site breached these thresholds for periods that were sustained for less than 2 weeks 4 times over the end of the experimental period and did lead to heat accumulation as degree heating weeks (Fig 3c).

### Traits associated with growth

*Acropora formosa* fragments that were transplanted to the shallows had a greater percentage change in relative bulk volume than those transplanted to the deep (Anova, *χ*^2^ = 33.913, d.f. = 1, *P* = <0.001, Fig 4a), irrespective of their location of origin. Location of origin had a separate significant effect on percentage change in relative bulk volume of fragments, those originating from the reef slope increased their bulk volume significantly more than those from the reef flat (Anova, *χ*^2^ =4.026, d.f. =1, *P* =0.045, Fig 4b).

**Fig 4.**
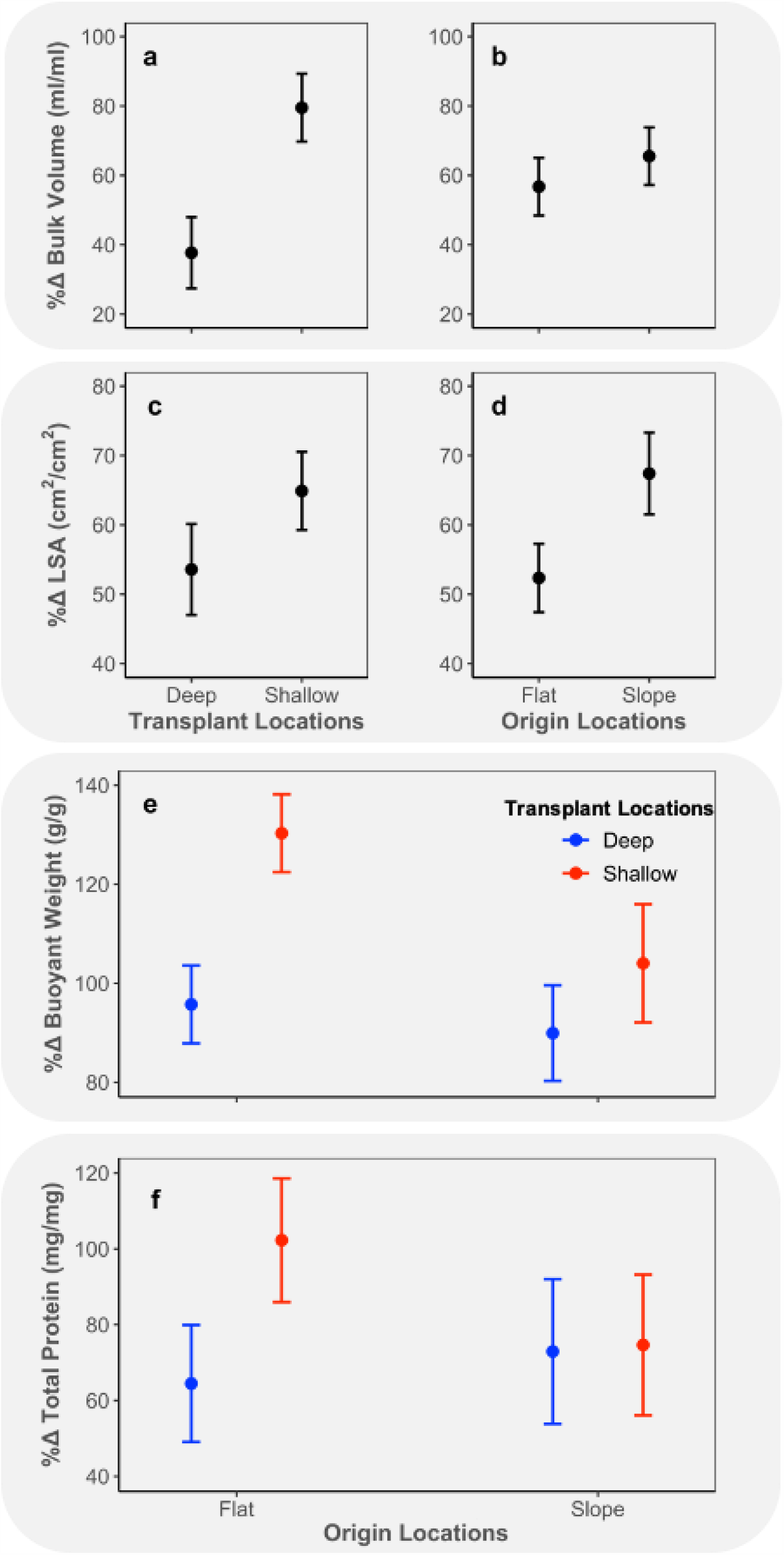
Change in bulk volume (ml/ml) of *A. formosa* fragments form Heron Island. (Southern GBR) by (a) location of Transplantation (P<0.05), and (b) location of Origin (P<0.05). Change in living surface area (LSA, cm^2^/cm^2^) of *A. formosa* fragments by (c) location of Transplantation (P<0.05), and (d) location of Origin (P<0.05) for fixed rate of change in volume. (e) Change in buoyant weight (g/g) for fixed changes in living surface area between location of Transplantation (shallow and deep), and location of Origin (flat and slope). (f) Change in total proteins (mg/mg) for fixed changes in living surface area and buoyant weight between location of Transplantation (shallow and deep), and and location of Origin (flat and slope). Error bars in all graphs represent a 95% confidence interval. All variables are reported as percentage relative change from initial over the experimental period, as indicated by “% Δ” in all axis titles. Each grey rectangle indicates figures derived from the same models (ie. a and b).

Analysis of relative rates of change in living surface area (LSA) found that when other parameters were held constant at mean value: (i) rates of change in LSA correlated positively with rates of change in coral volume (Anova, *χ*^2^ = 171.814, d.f. = 1, *P* = <0.001); (ii) rates of change in LSA were significantly greater in corals transplanted to reef flat compared to those transplanted to reef slope (Anova, *χ*^2^ = 5.601, d.f. = 1, *P* = 0.018, Fig 4c); (iii) rates of change in LSA were significantly greater in corals of reef-slope origin than in corals of reef-flat origin (Anova, *χ*^2^ = 14.708, d.f. =1, *P* = <0.001, Fig 4d). For the same rate of change in volume, changes in coral LSA were greater in reef slope origin corals and tended to indicate increased surface rugosity as opposed to reduced partial tissue mortality.

Relative rates of change in buoyant weight (BW) between *Acropora formosa* fragments analysed when other parameters were held constant at mean value found that: (i) rates of change in BW correlated positively with rates of change in fragment LSA (Anova, *χ*^2^ = 24.383, d.f.= 1, *P* = <0.001); (ii) corals that originated from the reef flat and were transplanted to the reef flat (“Flat/Shallow”) had significantly higher rates of change than any other group (Anova, *χ*^2^ = 4.980, d.f.= 1, *P* = 0.026, Fig 4e), which were not significantly different from each other.

Analysis of relative rates of change in total proteins found that when other parameters were held constant at mean value: (i) rates of change in total proteins correlated positively with rates of change in LSA (Anova, *χ*^2^ = 6.342, d.f.= 1, *P* = 0.012); (ii) rates of change in total proteins were significantly greater in fragments originating from the reef flat transplanted to the shallow, than transplanted to the deep (Anova, *χ*^2^ = 5.046, d.f.= 1, *P* = 0.025, Fig 4f), but not different between reef slope origin corals transplanted to the shallow or deep.

Relative rates of change in total lipids of *A. formosa* fragments when analysed with other parameters held constant at mean value were significantly correlated with LSA (Anova *χ*^2^ = 11.948, d.f. = 1, *P* = <0.001). But, showed no significant effect of fragment origin or transplant location.

## Discussion

Understanding the responses of coral reefs to climate change is an imperative if we are to devise effective management and policy responses to climate change. In this regard, the genetic adaptation of corals to rapidly changing conditions has generally been considered to be too slow to keep up with climate change (Hoegh-Guldberg 2012). Nonetheless, recent literature has identified potential solutions that depend on inhibiting the bleaching response of corals either via epigenetics or genetic manipulations (Oliver and Palumbi 2011; van Oppen et al. 2011), or, by reducing irradiance over selective periods that coincide with the advent of thermal stress (Jones et al. 2016). Observations of recent intense thermal events on the GBR, however, suggest that delaying the onset of bleaching in a thermal event provides no safeguard against coral mortality (Hughes et al. 2018a). Additional, it is also accepted that even increased coral survival rates will be insufficient to maintain the ecological function of coral reefs if they fail to equate with the maintenance of a net positive carbonate balances for the reef ecosystem (Andersson and Gledhill 2013; Perry et al. 2013). In the present study, we set out to test an alternative hypothesis that corals survive thermal stress because they conserve energy through reducing branch extension rates. Energy that can be redirected into the maintenance of existing tissue biomass. Investment in protein as opposed to skeletal growth leads to increased energy reserve capacities (Rodrigues and Grottoli 2007), repair and maintenance capacities and potentially even the coral’s heterotrophic capacity, which at minimum requires the production of digestive enzymes to breakdown particulates for uptake over the gastrodermis.

We hypothesized that long-term exposure to the highly variable reef-flat environment would both increase survival potential and engender a conservative growth response, where greater biomass is accumulated at the cost of conserving primary calcification rates. A conservative response that would be maintained upon transplantation to the new reef slope environment. The results of our study supported our hypotheses, and thereby, suggest that any potential epigenetic memory implanted by exposure to long-term prior stress is unlikely to be a panacea that will save carbonate coral reefs in the short-term. It is highly likely that coral survival increased because primary calcification is to some extent relinquished. In the present study, coral properties were maintained over 4 months. Other studies, however have demonstrated that epigenetic memories can remain in some organisms for generations (Klosin et al. 2017). Consequentially, this conservative response. has the potential to reduce carbonate coral reef resilience, the ability of carbonate reefs to bounce back from damage incurred from any disturbance events that engenders a loss of 3D framework or carbonate from the reef ecosystem (Connell 1997). The resilience of shallow tropical reefs may slide towards that of deep-cold water reefs that are able to establish as carbonate reefs, over tens of thousands of years, only because disturbance events are rare (presently, mostly man-made and occur in the form of drag-nets), and reef erosion rates scale with reef calcification rates (Kleypas et al. 1999b; Freiwald 2002; Bongaerts et al. 2010).

The environmental contrast between the reef slope and reef flat is exhibited by the difference between the mean and variability of the recorded PAR and temperature. The high variability in temperature on the reef flat observed in this study is comparable to temperature fluctuations observed in studies identifying specific coral populations with superior thermal tolerance (Oliver and Palumbi 2011; Howells et al. 2012). Degree heating weeks (DHW) products tend to use the hottest pixels to determine regional SSTs and MMMs that are based on these SSTs (Liu et al. 2014), and we mirrored this by using the third quartile (Q3) daily temperatures which gave rise to DHWs that accumulated to 8°C weeks over the height of summer, but only on the reef flat, not on the reef slope. Despite this apparent difference in heat stress, there was no apparent (visible) bleaching irrespective of coral origins (Fig 2), only significant mortality differences by coral Origin, but not experimental location.

Whilst mean temperature between the reef flat and reef slope did not differ significantly, mean PAR did. Based on this data, we conclude that the primary difference between the deeper slope and shallower flat zones is light. However, there could be additional confounding factors such as differences in the concentration of available dissolved or particulate organic matter that were not measured, but have been described in other studies (Done 1983; Wild et al. 2004; De Goeij et al. 2013). The higher light environment of the reef flat appeared to promote primary calcification, whilst experimental positioning on the deeper slope reduced all aspects of coral growth from primary and secondary calcification to protein accumulation. Nonetheless, reef slope origin corals experienced significantly greater primary calcification than reef flat origin corals at both experimental locations. Greater light and/or greater availability of organic nutriment may increase the energy available for growth processes to reef flat located corals relative to reef slope located corals over the long-term.

Over the experimental period of 4 months, reef flat origin corals showed a reduced ability relative to corals originating from the reef slope to acclimate to their new environment. Reef flat origin corals were unable to increase their energy acquisition to maintain the additional energy required to increase skeletal and protein densities on the slope, despite their general tendency to exhibit slow volume and tissue expansion rates. This could be, either a result of symbionts that do not acclimate to the new light environment, and/or because organic nutrients are less available at the interface between the ocean and the reef, than within a ponding reef-flat where excreted dissolved organic carbon can be enriched by microbial activity (De Goeij et al. 2013; Rix et al. 2017). By contrast, reef slope origin corals that expand relatively rapidly irrespective of location, were not apparently photo-inhibited or photo-oxidised (Richier et al. 2008) by the move to the higher light regime, but actually benefited from an apparent increase in available energy observed as a slight increase in rates of secondary calcification. This is contrary to expectations. The literature tends to argue that zooxanthellate corals can be photo-inhibited, potentially resultant from photooxidation, by increases in light, especially if light increases come in combination with increases in thermal stress (Richier et al. 2008; Skirving et al. 2018). But, many corals can acclimate over days to weeks to reduction in light (Anthony and Hoegh-Guldberg 2003; Cohen and Dubinsky 2015).

A potential reason the corals benefitted from the higher light environment in our study is that the high light habitat considered is potentially relatively richer in dissolved and particulate organic food (Wild et al. 2004; De Goeij et al. 2013). Corals with thicker tissues and reduced rugosities, have been linked to a reduced ability of dinoflagellate endosymbionts to respond to external light fluxes due to the probability that the skeleton plays a lesser role in increasing the path-length of these photons (Enríquez et al. 2005). Here, host pigment and other tissue structures compete with symbionts to absorb photons (Fisher et al. 2012; Pontasch et al. 2017). This could be the optimal response of a coral subjected to highly variable light dosing through time, that otherwise may consume significant energy as they rebuild light harvesting structures and repair D1-protein when light fluxes exceed the capacity of short-term responses such as xanthophyll cycling (Hill et al. 2011; Jeans et al. 2013). Such hosts may depend more on heterotrophy. The data from our study does not suggest that primary calcification on the reef flat is limited by the available ‘head space’ of water above colonies (Pandolfi et al. 2011). This experiment was performed on small fragments that were not limited by water head space at either location. Increasing water depth, is likely to reduce light levels unless corals have the capacity to keep up, and/or acclimate their energy acquisition in response to the reduction in light (Perry et al. 2018). Reef-flat origin corals in this study did not demonstrate this potential.

We have demonstrated here that conspecific of *A. formosa* that showed greater potential to survive a diversity of environments appear to exhibit a reduced ability to acclimate to changes in light availability. Attempting to further protect such corals from bleaching by periodically reducing light fields, by cloud seeding (Jones et al. 2016), would therefore impart further restrictions on energy acquisition and calcification. By contrast, the translocation of reef slope communities to the reef flat, did not generate any evidence of bleaching, despite the significant increase in the light field and regular incursions into stressful temperatures. Given that the mortality rate of these corals did not vary by location, the evidence suggests that Acroporids originating from the reef slope acclimate to the changing light regime relatively quickly (21 days; Anthony and Hoegh-Guldberg 2003; Jeans et al. 2013). Rapid photo-acclimation however has significant ramifications on the efficacy and practicality of applying shade to protect such corals should they be exposed to greater heat stress. Shade is intended to reduce excitation pressure at PSII and prevent the photo-oxidation to the Symbiodiniaceae photosystems, but if these photosystems rapidly increase the efficiency of photon capture in response to reductions in the light field, then shade will not serve to reduce excitation pressure. Shading will however consume additional energy as symbionts would need to invest in rebuilding their light harvesting antennae (Stambler and Dubinsky; Titlyanov et al. 2001), potentially reducing quanta of energy translocation to host.

On a more positive note, reef flat coral transferred back to the reef flat had higher skeletal density relative to other treatments, whilst also maximising protein density. Consequentially, primary calcification was reduced. Increases in skeletal density can reduce fragmentation and over the long-term (Lirman 2000; Okubo et al. 2007), even colonies with limited primary calcification can accumulate to cover a large area in space as long as physical forces do not exceed the strength of the produced skeleton (Scoffin et al. 1992). Notably lower expansion rates would be beneficial to the maintenance of symbiont densities, as the need for production of new symbionts to occupy the new material would be reduced. Rapid outward expansion is reportedly associated with “bleached” coral tissue (Oliver 1984). Clearly, some of these hypotheses regarding the explanation of the observed responses require testing. If epigenetic memory is responsible for increasing the survival of reef flat corals exposed to highly variable environments, then environments that tandem variability in light and temperature, in addition to many other abiotic variables, result in corals that sacrificed calcification (Denis et al. 2011, 2013). Survival appears not to be based on the reduced ability to bleach, but rather correlates with reductions in volume and tissue expansion in the Acroporid coral tested. Other corals (Gold and Palumbi 2018), may maintain rates of linear extension, but it is questionable whether other aspects of calcification are not compromised. For example, reef flat and reef slope conspecifics of *A. formosa* from this study, clipped to the same length, have different volumes and skeletal densities that forced the use of relative changes to preserve the statistical assumption that covariates are independent of treatment effect.

Presently, we assume that the differences observed between these relatively proximal *A. formosa* populations are driven by epigenetic consequences associated with prior long-term exposure to different abiotic environments within the reef-scape. There is, however, the potential that the relative inflexibility of the two coral populations to alter properties when exposed to new environments has a genetic cause. The two populations may hide cryptic speciation, a phenomenon that is increasingly observed amongst coral taxa (Rosser 2016; Gold and Palumbi 2018; van Oppen et al. 2018). For cryptic speciation to occur amongst spawning populations that are relatively proximal in space, gene flow may be limited (Nosil and Crespi 2004). In this regard, temperature can change the timing of spawning (Keith et al. 2016), and asynchronous spawning can inhibit gene flow (Rosser 2016). Alternatively, cryptic species could exhibit different larval and settlement properties that reinforce differential recruitment in these zones (Combosch and Vollmer 2011). In a brooding species, *Seriatopora hystrix* genetic isolation has been observed to result from differences in reproductive timing between populations in deep and shallow habitats (Bongaerts et al. 2011, 2020). More recently the same has been determined for *Pocillopora damicornis* (van Oppen et al. 2018). Interestingly, this genetic differentiation in *P. damicornis* was observed on Heron Reef over similar geographic distances as in the current study. However, *P. damicornis* is a brooding species while the target species in the current study, *A. formosa*, is a broadcasting species.

Corals that have a prior life history of chronic exposure to highly variable environments tend to be less susceptible to mortality. In this study, we observed that corals raised to adult colonies on the reef flat have a greater survival potential in any environment than conspecifics raised on the reef slope. Daily third quartile temperature accumulated to 8°Cweek over the summer on the reef flat. In combination with an increased irradiance for reef slope origin corals moved to the reef flat, the new environment however did not increase bleaching susceptibility of these reef slope origin corals. Rather, they demonstrated that coral expansion in these slope origin corals was suppressed by the lower light conditions under which they appear to nonetheless flourish on the reef slope. The greater survival potential of reef flat origin corals was correlated to their having greater tissue thickness likely facilitated by reduced rates of primary and secondary calcification. Furthermore, these reef flat origin corals failed to acclimate to the lower light levels offered on the reef slope as suggested by a lack of energy to maintain high tissue protein densities despite reduced branch expansion. The fact that reef flat origin corals do not physiologically morph into reef slope origin corals upon relocation to the reef slope and vice-versa suggests that these corals have become epigenetically or genetically distinct populations. This study supports the hypothesis that exposure to a highly variable environment results in “tough” corals, but it identifies that one of the costs of increased survival potential is reduced calcification. These “super corals” therefore lack the properties that are required to save carbonate reefs exposed to increases in the intensity and frequency of disturbance events as a result of climate change. Further, they lack the ability to respond positively to the reductions in light field likely associated with future sea-level rise. Branching Acroporids are important to the 3-dimensional structure and many services provided by carbonate reefs. The present study suggests that the characteristic that allow corals to survive and prosper in highly variable reef environments, do not align with those that are required to maintain carbonate coral reefs exposed to the many consequences of climate change.

## Acknowledgements

We thank the members of the Coral Reef Ecosystems laboratory at UQ, especially Aaron Chai and Josh Biggs, for assistance with coral collection and preparation. Draft reviews by Ove Hoegh-Guldberg, Catherine Kim, and Kristen Brown improved the manuscript. The experiment was funded by the Australian Research Council (ARC) and by the National Oceanic and Atmospheric Administration (NOAA) via ARC Centre of Excellence for Coral Reef Studies CE140100020 (S.D and O.H-G) and ARC Linkage LP110200874 (O.H-G and S.D.).

